# Urinary proteome changes during pregnancy in rats

**DOI:** 10.1101/2022.08.04.502874

**Authors:** Shuxuan Tang, Youhe Gao

**Affiliations:** Gene Engineering Drug and Biotechnology Beijing Key Laboratory, Life Science College, Beijing Normal University, Beijing 100875, China

**Keywords:** Urinary proteome, Normal pregnancy, Embryonic development, Coagulation pathway

## Abstract

Pregnancy involves a significant amount of physiological changes. A normal pregnancy is essential to ensure healthy maternal and fetal development. We sought to explore whether the urinary proteome could reflect the pregnancy process. Urine samples were collected from pregnant rats on gestational day 1, 4, 7, 11, 14, 16, 18, 20 (GD 1 d, GD 4 d, GD 7 d, GD 11 d, GD 14 d, GD 16 d, GD 18 d, GD 20 d), and control rats on days 0, 4, 7, 11, 14, 16, 18 and 20. The urinary proteome was profiled by liquid chromatography coupled with tandem mass spectrometry (LC-MS/MS), and differential proteins were obtained by comparing the 0 d (GD 1 d) of the same group at each time point within the two groups. Through the analysis of the enriched pathways of differentially expressed proteins in the pregnant group, during the period from fertilization to implantation, many pathways related to embryo implantation and trophoblast differentiation were enriched on GD 1 d, GD 4 d and GD 7 d. In addition, the developmental process of the fetal rat heart such as heart looping and endocardial cushion formation, are consistent with the timing of previous studies; the developmental process of the lung and the development of the rat embryo alveoli before birth are consistent with the reported timing; and the developmental time of the rat embryo pancreas is also during the period of pancreatic cell proliferation and differentiation. These processes were enriched only in the pregnancy group and not in the control group. Furthermore, coagulation-associated pathways were found to be increasingly prominent before labor, which is consistent with the previously reported trend of increasing coagulation function during pregnancy. Our results indicated urinary proteome can reflect some embryonic developmental and maternal changes in rat pregnancy.

## 1 Introduction

Pregnancy is a process of great physiological changes in the body, including numerous metabolic and hormonal adaptations (1). Pregnancy proceeds through placental development, angiogenesis, remodeling of uterine tissue, and inflammation, resulting in the release of various hormones and cytokines (2). A small error in the normal pregnancy process can lead to serious consequences. For example, disruption of certain pregnancy processes can lead to cervical dysfunction, infection and spontaneous preterm rupture of membranes and preterm labor (2). How to better ensure a healthy pregnancy is very important for the normal development of the fetus and reducing the damage to the mother.

Ultrasound is currently used clinically as a method of monitoring fetal development in combination with chorionic villus sampling, amniocentesis, cortical puncture, and the determination of specific maternal plasma markers of abnormal fetal abnormalities (3).

Suitable maternal biological fluids (e.g., urine and blood) are used to characterize the state of a pregnancy, and ensuring normal fetal development over time is critical. With the development of high-throughput technology, proteomics technology provides a new strategy for the discovery of biomarkers with high-throughput and sensitive characteristics, and omics is increasingly widely applied in disease diagnosis and treatment monitoring. The urinary proteome is a noninvasive assay used to detect maternal abnormalities and embryonic diseases during pregnancy, such as preeclampsia (4,5), gestational diabetes (6), gestational trophoblastic disease (7), gestational hypertension (8), trisomy 21 (9), and aneuploidy (10).

Urine is easier to collect than other body fluids, urine is not regulated by body homeostasis, so urine can reveal early disease-related changes, earlier than blood indicators (11). The urinary proteome reflects changes in different types of diseases, such as Parkinson’s disease (PD) (12), autism (13), glioma (14), liver cancer (15), bone cancer metastasis (16), colitis (17), and myocarditis (18). In this study, we explored whether the urinary proteome could reflect maternal changes and the process of embryonic growth and development through animal experiments.

In this study, urine was collected on GD 1 d, GD 4 d, GD 7 d, GD 11 d, GD 14 d, GD 16 d, GD 18 d, and GD 20 d in the pregnant group. In the control group, the urine of nonpregnant rats was collected on the corresponding 0 d, 4 d, 7 d, 11 d, 14 d, 16 d, 18 d, and 20 d, and the urinary proteome was analyzed by high-throughput LC-MS. The quantitative results of the urinary proteome at each time point in the pregnant group were compared with those at GD 1 d. Correspondingly, the quantitative urinary proteome at each time point in the control group was compared with that at 0 d to determine the differential proteins. The results showed that the urinary proteome could reflect changes in different stages of pregnancy, including maternal physiological changes and embryonic development-related processes. The study design flowchart is presented in Fig. 1.

**Fig. 1.**
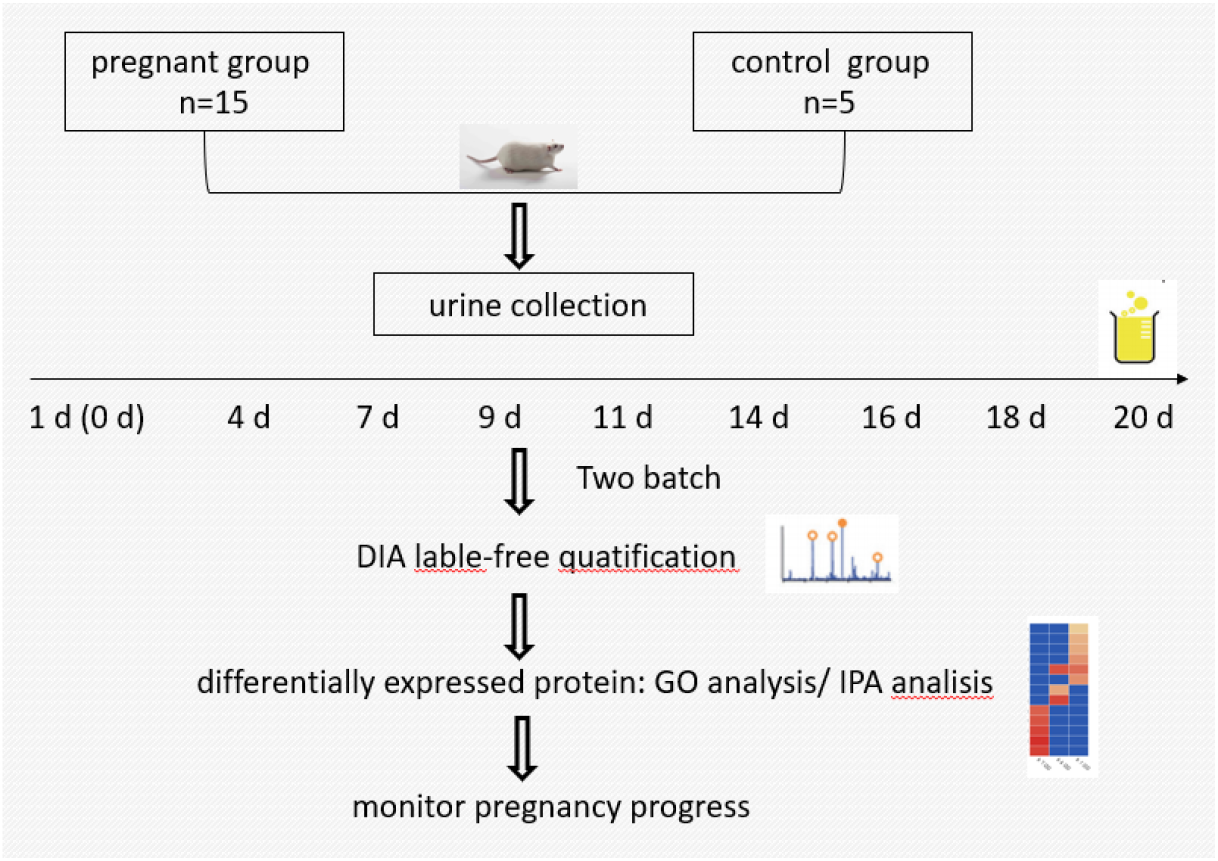
Flowchart of the study design. DIA, data-independent acquisition; biological processes analysis of differential proteins was performed by GO

## 2 Materials and methods

### 2.1 Experimental animals and groups

Pregnant Wistar rats (GD 1 d, n=15, 190±40 g) and female Wistar rats (n=5, 200 ±40 g) were purchased from Beijing Vital River Laboratory Animal Technology Co., Ltd. The day on which the vaginal plug of the female rat was examined was GD 1 d. The animal room was maintained at a temperature of 25±2 °C, a humidity of 65-70%, and a 12-hour light-dark cycle. This study was conducted in accordance with guidelines developed by the Institutional Animal Care and Use Committee of Peking Union Medical College. This experiment was approved by the Ethics and Animal Welfare Committee of Beijing Normal University (CLS-EAW-2020-022).

In this study, Wistar rats were divided into two groups: pregnant group and control group. The urine samples of the two groups were collected in two batches. The first batch of urine samples was collected from pregnant rats (n=10): GD 1d, GD 14 d, GD 16 d, GD 18 d and GD 20 d. The second batch of urine samples was collected from pregnant rats (n=5) and control rats (n=5): pregnant group: GD 1 d, GD 4 d, GD 7 d, GD 11 d; control group 1 d, 4 d, 7 d, 11 d, 14 d, 16 d, 18 d, 20 d.

### 2.2 Urine sample preparation

Urine was collected from the pregnancy and control groups on Days 1, 4, 7, 11, 14, 16, 18, and 20. Urine (4-6 ml) was centrifuged at 14000*g for 30 min at 4 °C. The supernatant was added to at least 3 volumes of ethanol. The solution was placed at −20 °C overnight for precipitation. Then, 100 μl of lysis buffer (8 mol/L urea, 2 mol/L thiourea, 50 mmol/L Tris, and 25 mmol/L dithiothreitol) was added. Protein concentrations were quantified by the Bradford assay.

Each sample was digested with 100 μg of protein using the FASP method (19). The sample was loaded onto a 10-kD ultracentrifugation filter and washed twice with UA solution (8 mol/L urea, 0.1 mol/L Tris-HCl, pH 8.5), and the protein sample was reduced with 20 mmol/L dithiothreitol (37°C) for 1 hour and then alkylated with 50 mmol/L iodoacetamide (IAA, Sigma) for 45 min in the dark. Next, the samples were washed twice with UA and three times with 25 mM NH4HCO3 and then digested overnight at 37 °C with trypsin (ratio of enzyme:protein 1:50). The digested peptides were eluted from 10-kD filters and desalted using HLB (Waters, Milford, MA, USA).

### 2.3 Peptide fractionation

The peptides were diluted to 0.0.5 μg/μl with 1%_0_ formic acid water, and all samples (1 μg/sample) were mixed into QC samples. The pooled sample was loaded on high pH reversed-phase spin columns (Thermo Scientific, 84868). Eluents with different concentrations of acetonitrile (5, 7.5, 10, 12.5, 15, 17.5, 20 and 50%) were added to reversed-phase fractionation spin columns, and the peptides were eluted with different concentrations of acetonitrile to obtain 8 different gradients of fractions. These peptides were lyophilized in vacuo and reconstituted in 20 μl of 1% formic acid in water.

### 2.4 LC-MS/MS analysis

To correct the retention time of the extracted peaks, iRT was added to each sample according to the ratio of peptides:iRT volume ratio of 10:1. One microgram of each sample was loaded at a rate of 300 μl/min through a trap column (75 μm * 2 cm, 3 μm, C18, 100 Å). Peptides were eluted with a gradient of 4%-35% buffer B (0.1% formic acid in 80% acetonitrile, flow rate at 0.3 μl/min) for 90 min.

Eight fractions were analyzed with a mass spectrometer in DDA mode. The parameters of MS in DDA mode were set as follows: the full MS scan was acquired from 350-1550 m/z with an orbitrap resolution of 12000. The MS/MS scan was acquired in orbitrap mode with a resolution of 30000. The HCD collision energy was set to 30%. The AGC target was set to 5e4, and the maximum injection time was 45 ms. Then, the mass spectrometer was changed to the DIA mode, and the individual samples were analyzed in DIA-MS mode. The variable isolation window of the DIA method with 36 windows was adopted. The full scan was set at a resolution of 60,000 over a range of 350-15000 m/z. All samples were mixed to obtain QC for DIA mode analysis.

### 2.5 Data processing

The ten raw files were searched against the Swiss-Prot rat database with SEQUEST HT by Proteome Discoverer (version 2.1; Thermo Scientific). A maximum of two missed cleavage sites were allowed in the trypsin digestion. The parent ion mass tolerances were set to 10 ppm, and fragment ion mass tolerance was set to 0.02 Da. The cutoff at the precursor and protein levels was less than 1%.

The QC and individual files were imported into Spectronaut Pulsar X. Protein inference was performed using the implemented IDPicker algorithm (20). The false Q value (FDR) was set at less than 1% for proteins and precursors.

### 2.6 Statistical analysis

Quantitative results were processed by Spectronaut. QC samples were used to evaluate the stability of the mass spectrometry. The sequential-KNN method was used to fill in the missing values of QC (https://www.omicssolution.org/wkomics/main). Proteins with a coefficient of variation of QC < 0.3 were selected as differential proteins for subsequent screening.

One-way ANOVA was used for the comparison of data at different time points. The proteome at each time point in the pregnant and control groups was compared with the 0 d (GD 1 d) proteome. Differential protein screening criteria: at least two specific peptides in the protein were identified (fold change ≥ 1.5 or ≤ 0.67, t test P < 0.5). All results are presented as the mean ± standard deviation (SD).

### 2.7 Bioinformatics and functional analysis

Biological processes (BPs) were obtained by GO analysis of the differentially expressed proteins using the DAVID website. Biological pathway analysis and disease/biological function analysis were performed with differential proteins by IPA software (Ingenuity Systems, Mountain View, CA, USA). Visual mapping and data preprocessing were conducted on the Wu Kong platform (https://www.omicsolution.org/wkomics/main/).

## 3 Results and discussion

### 3.1 Urine proteome changes and function analysis in pregnant and the control group

In this study, urine samples from the rats in the pregnant group (n=15) and the control group (n=5) were collected at GD 1 d, GD 4 d, GD 7 d, GD 11 d, GD 14 d, GD 16 d, GD 18 d, GD 20 d and 0 d, 4 d, 7 d, 11 d, 14 d, 16 d, 18 d, 20 d. The number of urine proteomes identified in the two groups is shown in Table 1. According to the screening standard FDR<1.0%, each protein contained at least 2 unique peptides. First, the missing value of the mix was filled through the Wukong platform, the CV value of QC was evaluated, and the protein with CV<0.3 was selected. The number of credible proteins obtained from the first batch and the second batch were 1199 and 1289, respectively (table 1).

**Table 1.**
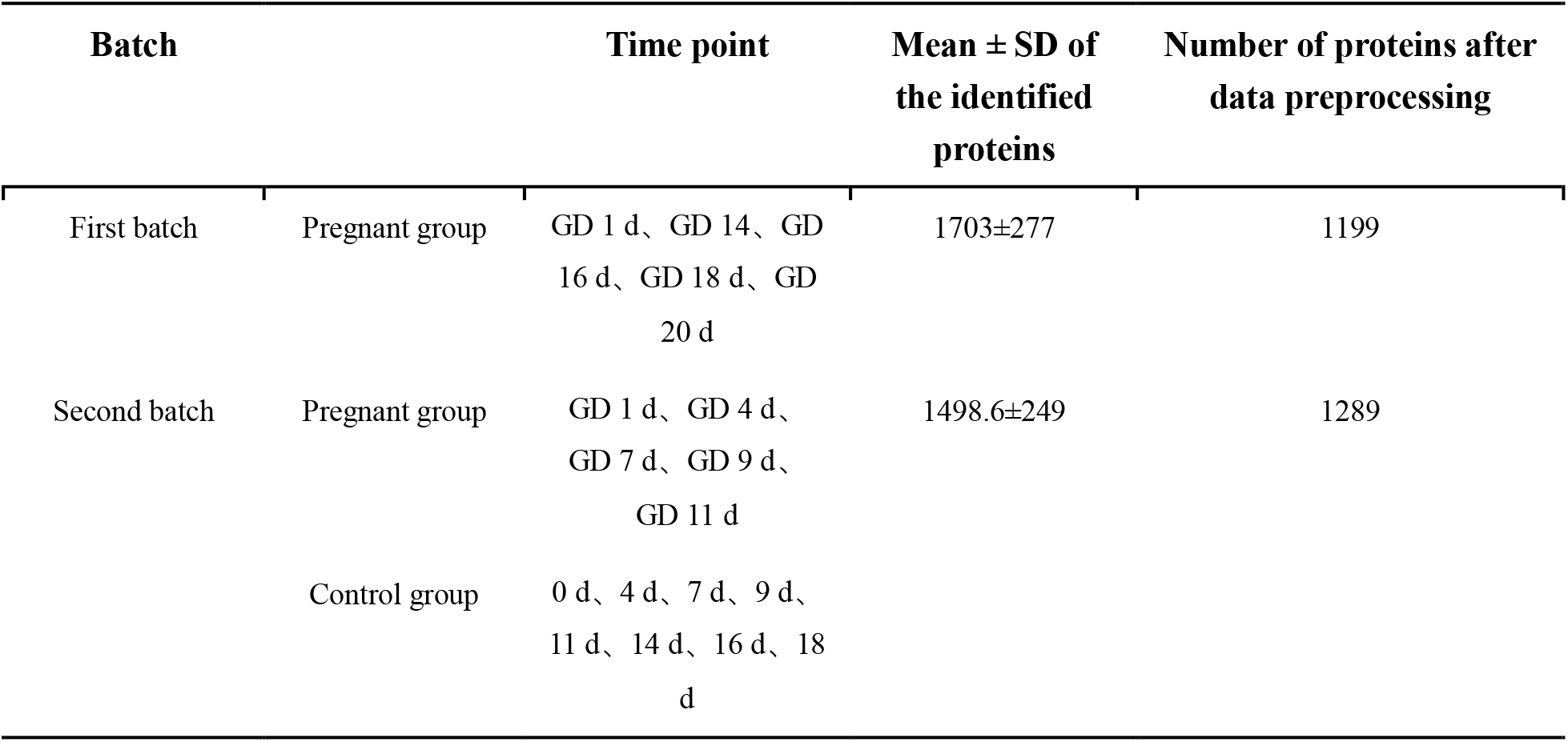
Urine proteome quantitative results

The differential proteins of the pregnancy group at each time point were obtained by comparison with the urine proteome of the same batch at GD 1 d. The differential proteins of the control group were obtained by comparison with the urine proteome at 0 d at each time point. A Venn diagram of differential proteins in the control and pregnancy groups is presented in Fig. 1A. There were 83 proteins in common between the control and pregnancy groups and 423 unique proteins in the pregnancy group, which were much more than those in the control group. We aimed to explore the physiological changes related to pregnancy from the urinary proteome, and the differential proteins of the pregnancy and control groups were enriched for biological processes and IPA biological classical pathways (Supplementary Table 1 & Table 2), respectively. The differentially expressed proteins of the pregnancy and control groups were enriched for biological processes and IPA biological classical pathways, respectively. The VEEN diagrams of the enriched results are shown in Fig. 1B and Fig. 11C. Based on a comparison of the number of differentially expressed proteins (Fig. 1A), biological processes (Fig. 11B) and IPA biological classical pathways (Fig. 1C) between the control and pregnant groups, the number in the pregnancy group was found to be much larger than that in the control group, suggesting that the changes in pregnancy reflected in urine were much greater than the changes associated with the growth and development of the rats themselves.

**Fig. 1a.**
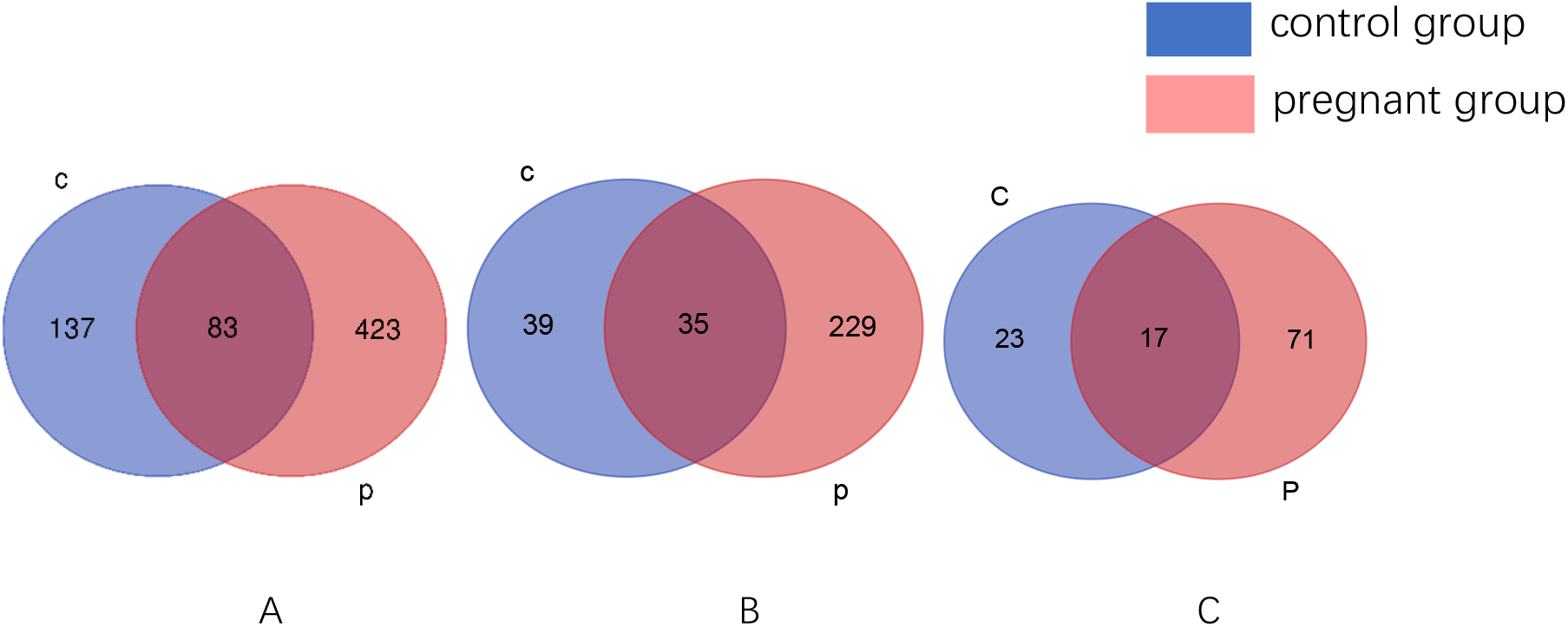
Differentially expressed proteins at all time points in the control and pregnant groups (A); biological processes at all time points in the two groups (B); IPA classical biological pathways at all time points in both groups (C).

### 3.2 Urine proteome changes during pregnancy

The differential proteins at each time point were identified and compared with those on GD 1 d. The urinary proteome changes on the 1st day after fertilization were reflected by the comparison of the urine proteome of GD 1 d and the control group at 0 d. After screening (FC≥1.5 or ≤0.67, p<0.05), 64, 56, 39, 69, 52, 69, 220, and 220 differential proteins were obtained at these time points in the pregnant group. A Venn diagram of the differentially expressed proteins in the pregnant group at each time point is shown in Figure 2. GD 18 d and GD 20 d had the largest number of differential proteins, which indicated that pregnancy changed greatly over the study time period.

**Fig. 2.**
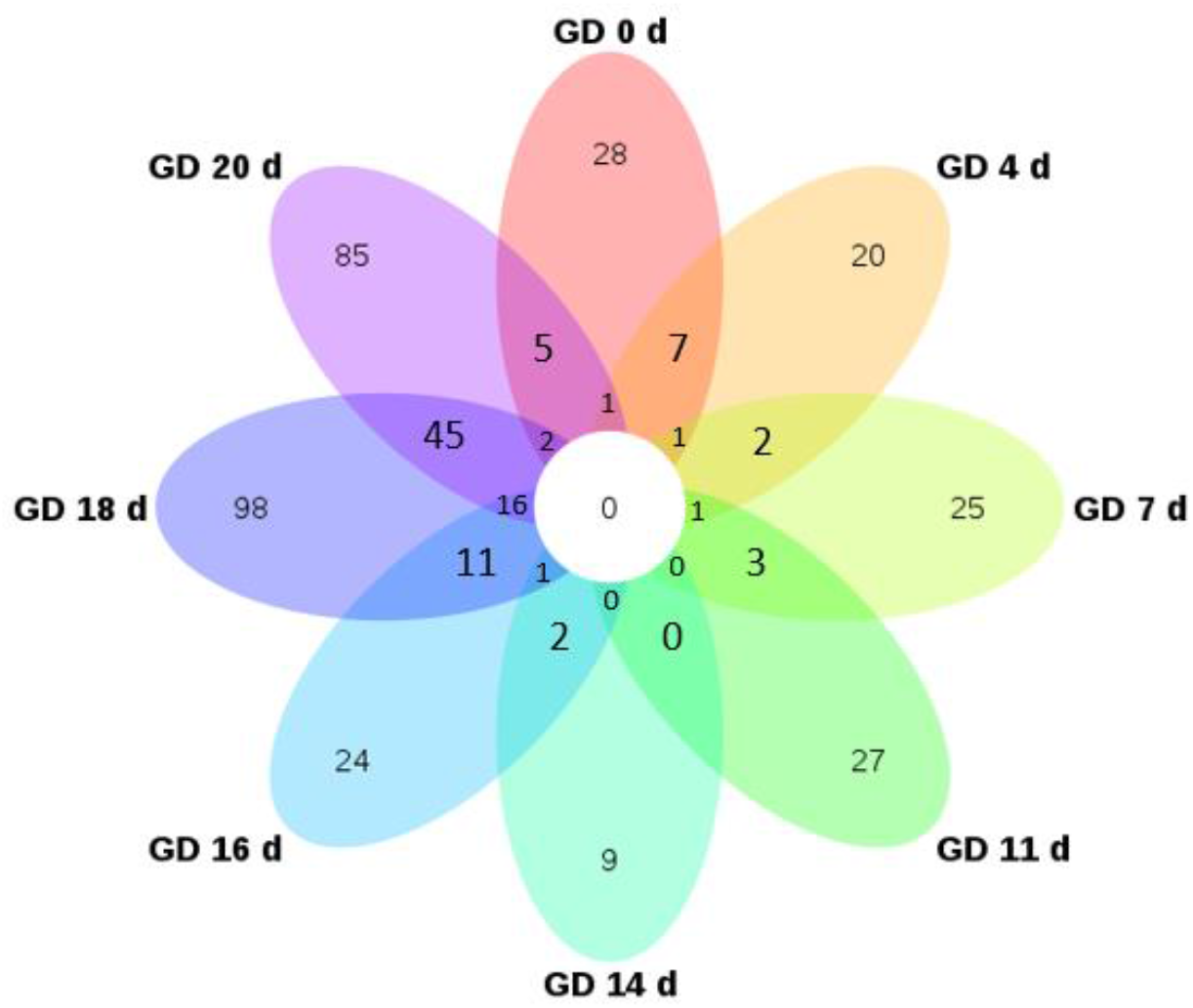
Venn diagram of differentially expressed proteins in the pregnant group

### 3.4 Functional analysis of the differentially expressed proteins in the pregnant group - implantation period

The implantation period of the rats started on GD 5 d and was completed on GD 7 d. The biological process of urine proteome enrichment of pregnant rats on GD 1 d, GD 4 d, and GD 7 d is shown in Figure 3 and Supplementary Table 3. Biological processes related to growth and development were enriched on the first day after fertilization (GD 1 d), including liver development, kidney development, and bone development, suggesting that proteins related to fetal growth began to work at a very early stage (Fig.3). Other biological pathways are the interleukin-17-mediated signaling pathway and response to L-ascorbic acid. Interleukin-17 activates the PPAR-γ/RXR-α signaling pathway and promotes the proliferation, migration and invasion of trophoblast cells (21). L-ascorbic acid (VitC) inhibits placental trophoblast apoptosis and increases syncytiotrophoblast differentiation, and VitC is also involved in the synthesis of placental steroids and polypeptide hormones (22).

**Fig. 3.**
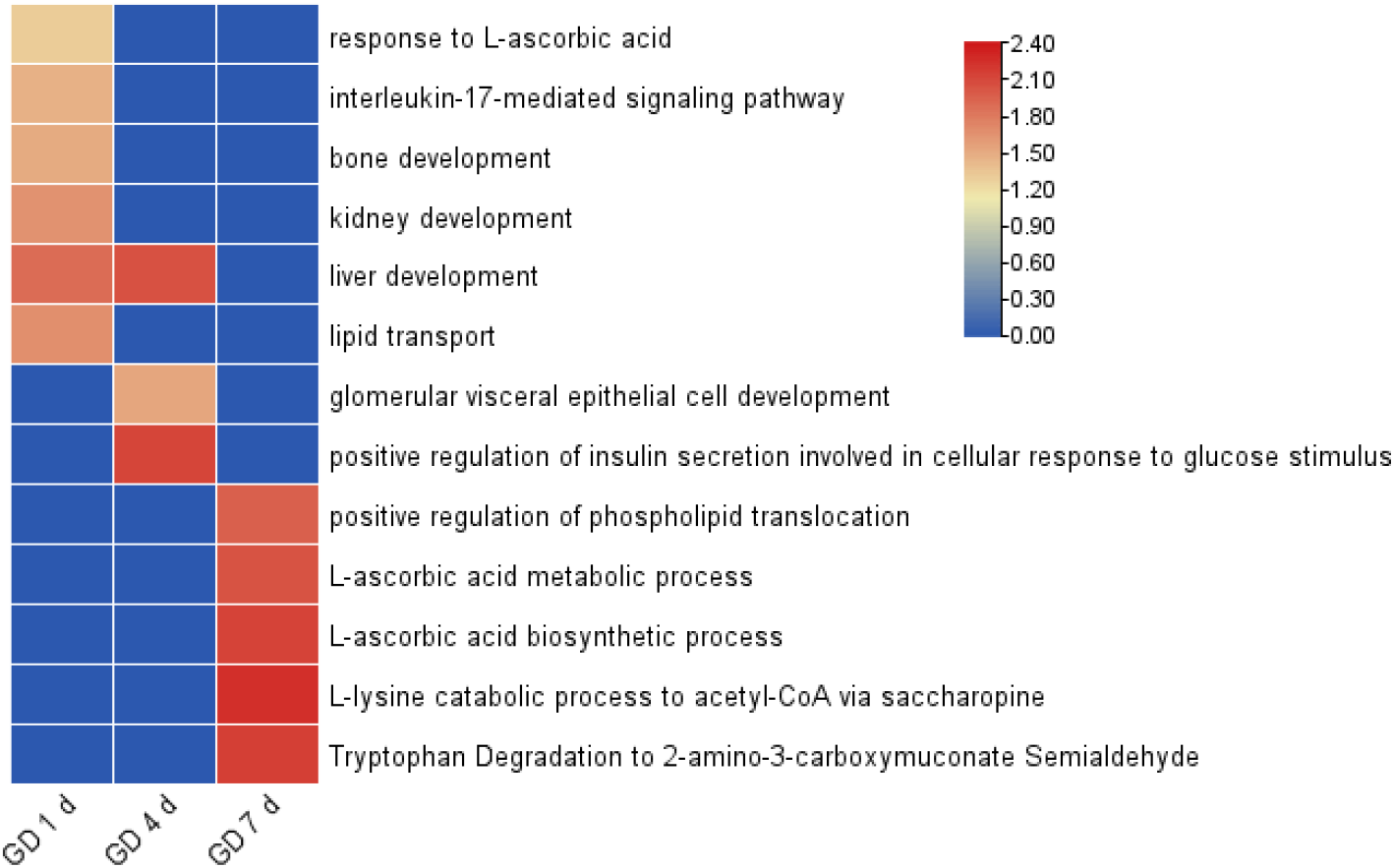
Biological pathways of differentially expressed proteins on GD 1d, GD 4 d and GD 7d by GO analyzed.

The growth- and development-related biological pathways displayed by differential proteins on GD 4 d of pregnancy include liver development and glomerular visceral epithelial cell development. Pathways affecting implantation include positive regulation of insulin secretion involved in the cellular response to glucose stimulus (Fig. 4). Insulin can directly impair ovarian function during embryo implantation and insulin-induced ovarian autophagy imbalance.

**Fig. 4.**
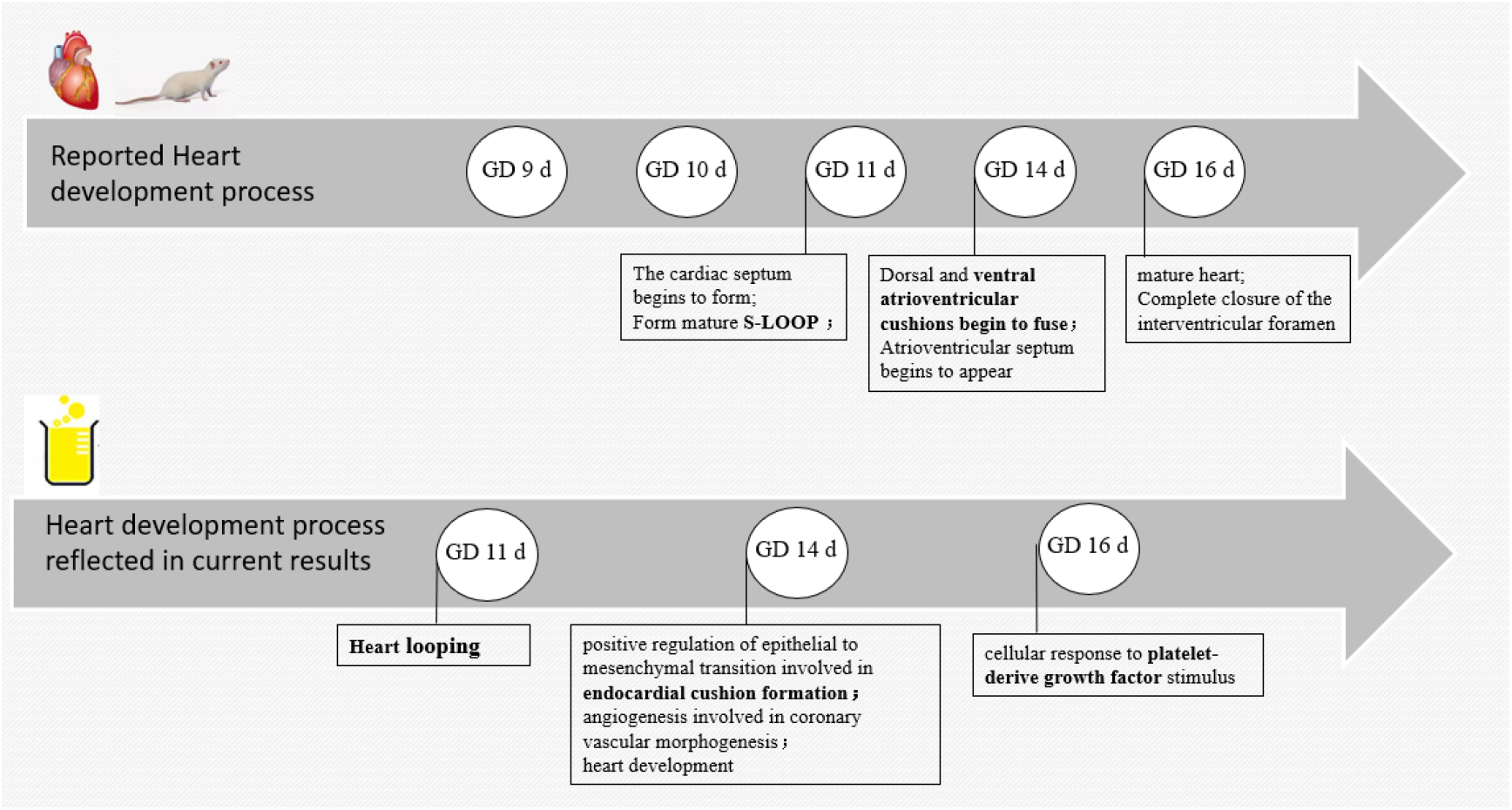
Fetal rat heart development The above is the reported heart development process; The following are caradiac developmental processes enriched by changing urinary proteomes

Many biological pathways and processes are related to implantation on GD 7d, such as the L-ascorbic acid metabolic process, L-ascorbic acid biosynthetic process and tryptophan degradation to 2-amino-3-carboxymuconate semialdehyde. Large amounts of ascorbic acid exert anti-inflammatory and immunological effects, which are beneficial for embryo implantation (23). Tryptophan is a key regulator of embryogenesis and embryo implantation during pregnancy. High L-Trp levels (500 and 1000 mM) induce cell cycle arrest in G1 phase and inhibit cell proliferation in porcine trophectoderm cells (24).

The urinary proteome exhibited during the implantation period of rats were mainly related to growth and development, implantation and placental growth, and the implantation-related pathways were only enriched in this period and did not appear in other periods the process of embryo implantation.

### 3.4 Functional analysis of the differentially expressed proteins in the pregnant group - embryonic organ development

Embryonic organ development is an important event in the process of embryo maturation. Rat embryonic organs begin to develop in the middle and late stages of pregnancy. The processes and pathways enriched by changes in the urinary proteome were analyzed to explore the growth and development of embryos.

Previous studies have shown that rat embryos begin to form a cardiac septum on GD 11 d and form a mature s-loop; on GD 14 d, the dorsal and ventral atrioventricular cushions begin to fuse, and the atrioventricular septum begins to appear; on GD 16 d, the heart matures, and the interventricular orifice is completely closed. The developmental period of the rat heart was reported to be from GD 9 d to GD 16 d (25). The development of the heart was reflected in the urine, as shown in Fig. 4. The study showed the biological process of heart looping on GD 11 d, the process of positive regulation of epithelial to mesenchymal transition involved in endocardial cushion formation on GD 14 d, and the cellular response to platelet–derived growth factor stimulus (PDGF) on GD 16 d. Clearly, the biological processes that emerged at GD 11 d and GD 14 d were completely consistent with the previously reported time course of cardiac development. PDGF-BB increased myocardium and myofibril compaction, and PDGF-AA increased myocardial fiber differentiation (26). In addition, angiogenesis involved in coronary vascular morphogenesis and heart development pathways was associated with cardiac coronary development.

Previous studies have shown that the pancreas of the rat developed a pancreatic bud at GD 9.5 d; the mesenchyme pushed the bulge around the dorsal gut to initiate pancreatic formation at GD 11 d; the mesenchyme surrounded the dorsal gut, pushing the bulge to initiate the formation of the pancreas at GD11 d-GD 11.5 d; and GD 14 d - GD 18.5 d is a critical period for rat pancreatic cell proliferation, differentiation and structure formation(27, 28). The present study shows that the urinary proteome reflects the process of pancreatic B cell proliferation at GD 14 d (Fig. 5), which is consistent with the previously reported period of pancreatic cell proliferation.

**Fig. 5.**
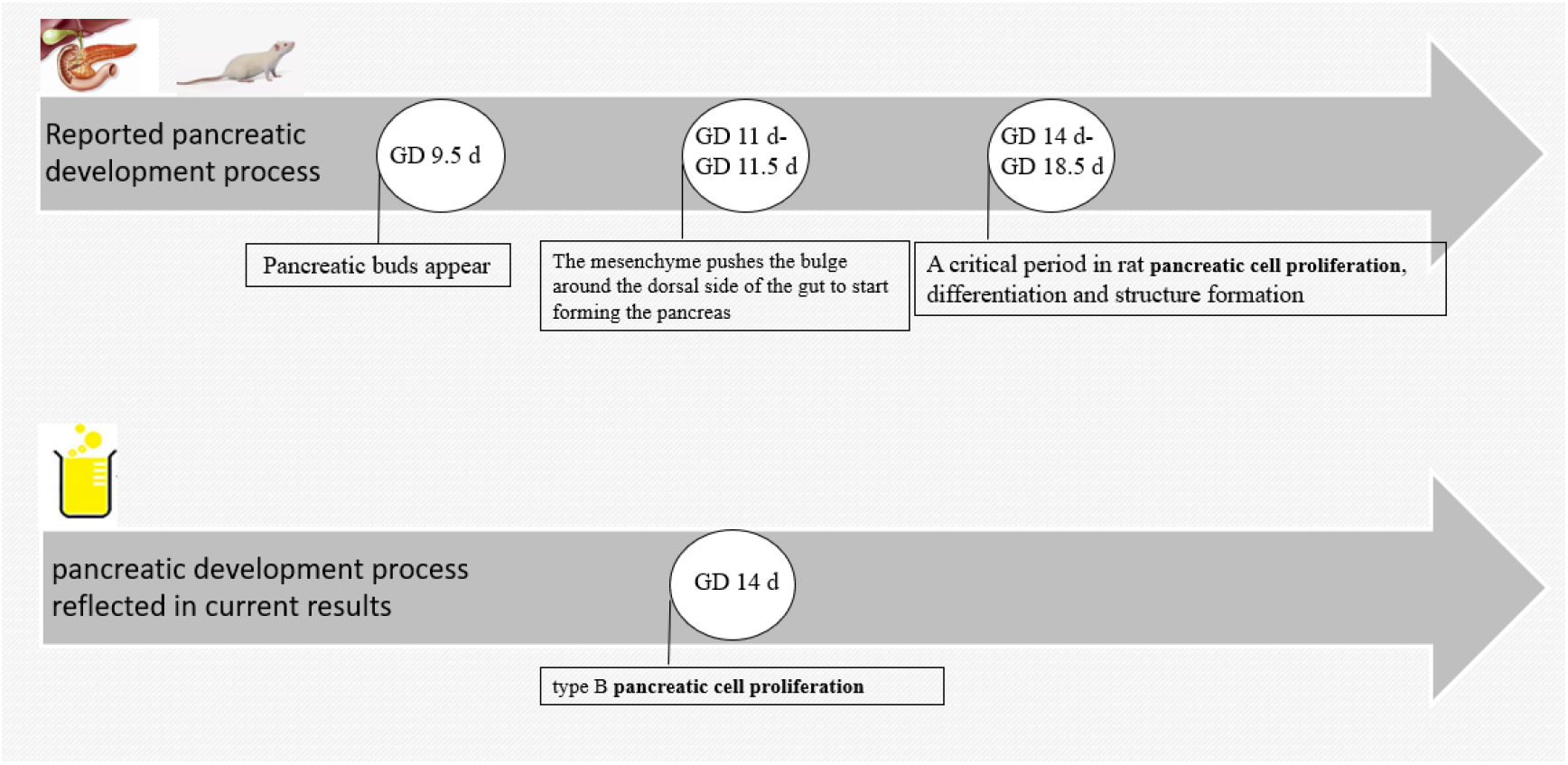
Fetal rat pancreas development. The above is the reported pancreatic development process. The following are pancreatic developmental processes enriched by changing urinary proteomes.

The development of embryonic lungs is closely related to the survival rate of the fetus after birth. The development of the rat embryonic lung begins on GD 11 d, and the lung primordium appears on GD 11 d; airway and pleura form on GD 13 d; bronchial development begins on GD 14 d; pulmonary alveoli and pulmonary vasculature generate between GD 14 d and 16 d, and alveolar and alveolar surfactant formation between GD 18.5 d and GD 20d (29). The results of this study showed that branching morphogenesis of an epithelial tube appeared on GD 11 d, the epidermal growth factor receptor signaling pathway (EGF) appeared on GD 16 d, and glucocorticoid receptor signaling, the Wnt/β-catenin pathway and PCP (planar cell polarity) appeared on GD 18 d (Fig. 6). The pathways emerging on GD 18 d are associated with alveolar and pulmonary surfactant formation, such as glucocorticoid Receptor Signaling, which increase dramatically in fetal plasma in late pregnancy, stimulating fetal pulmonary surfactant synthesis (30). Wnt/β-catenin is associated with airway smooth muscle cell proliferation and alveolar development (31). Alveolar formation during this period is approximately the same as previously reported. PCP is involved in the directional movement of cilia in the lungs (32). EGF at GD 16 d is an important regulator of cell differentiation and fetal pulmonary surfactant synthesis (33), possibly in preparation for the next stage of alveolar synthesis. Similarly, the biological process on GD 11 d may be preparation for the bronchial branches of the fetal lung, suggesting urine may reflect the developmental stage of the lung earlier during embryonic lung development.

**Fig. 6.**
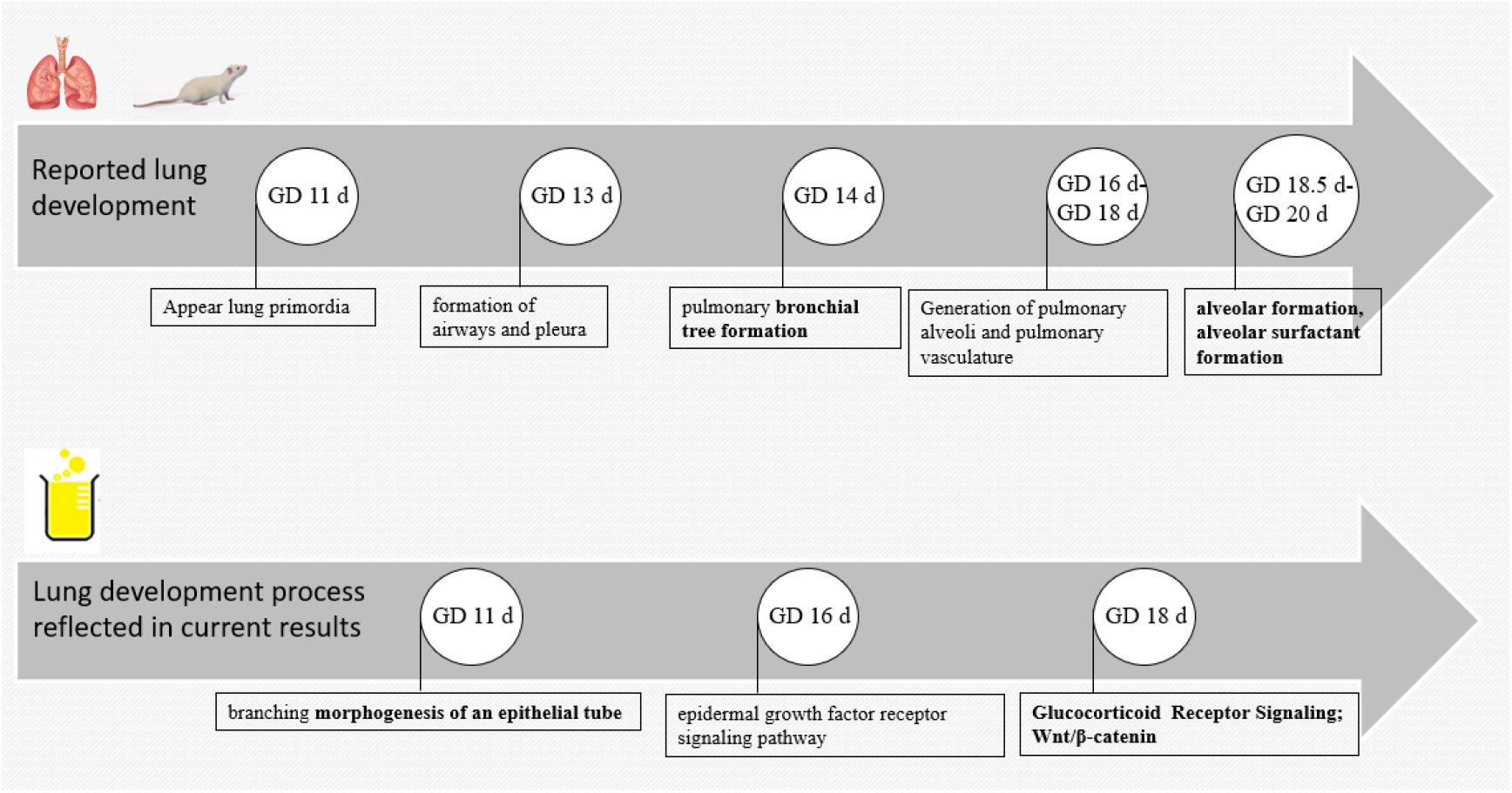
Fetal rat lung development. The above is the reported lung development process. The following are lung developmental processes enriched by changing urinary proteomes.

These pathways that corresponded to the timing of organ development only appeared in the pregnancy group and were not found in the control group (Supplementary Table 2), indicating that these pathways originated from embryonic development rather than the rat’s own growth and development. In addition, the process of growth and development of other organs over time is shown in Fig. 7. The pathways related to kidney development were glomerular visceral epithelial cell development and glomerular filtration. These two pathways were most prominent on GD 20 d, and glomerular visceral epithelial cell development was enriched on GD 4 d, GD 11 d, GD 16 d and GD 20 d; glomerular filtration was enriched on GD 16 d and GD 20 d; and liver development was enriched on GD 0 d, GD 4 d, and GD 11 d. Hair follicle morphogenesis was enriched on GD 11 d; salivary gland cavitation was enriched on GD 16 d; and skeletal muscle tissue development and neural crest cell migration were enriched on GD 18 d (Supplementary Table 4). These results suggest that the urinary proteome may reflect the status of embryonic growth and has the potential to be an effective strategy for monitoring normal embryonic development.

**Fig. 7.**
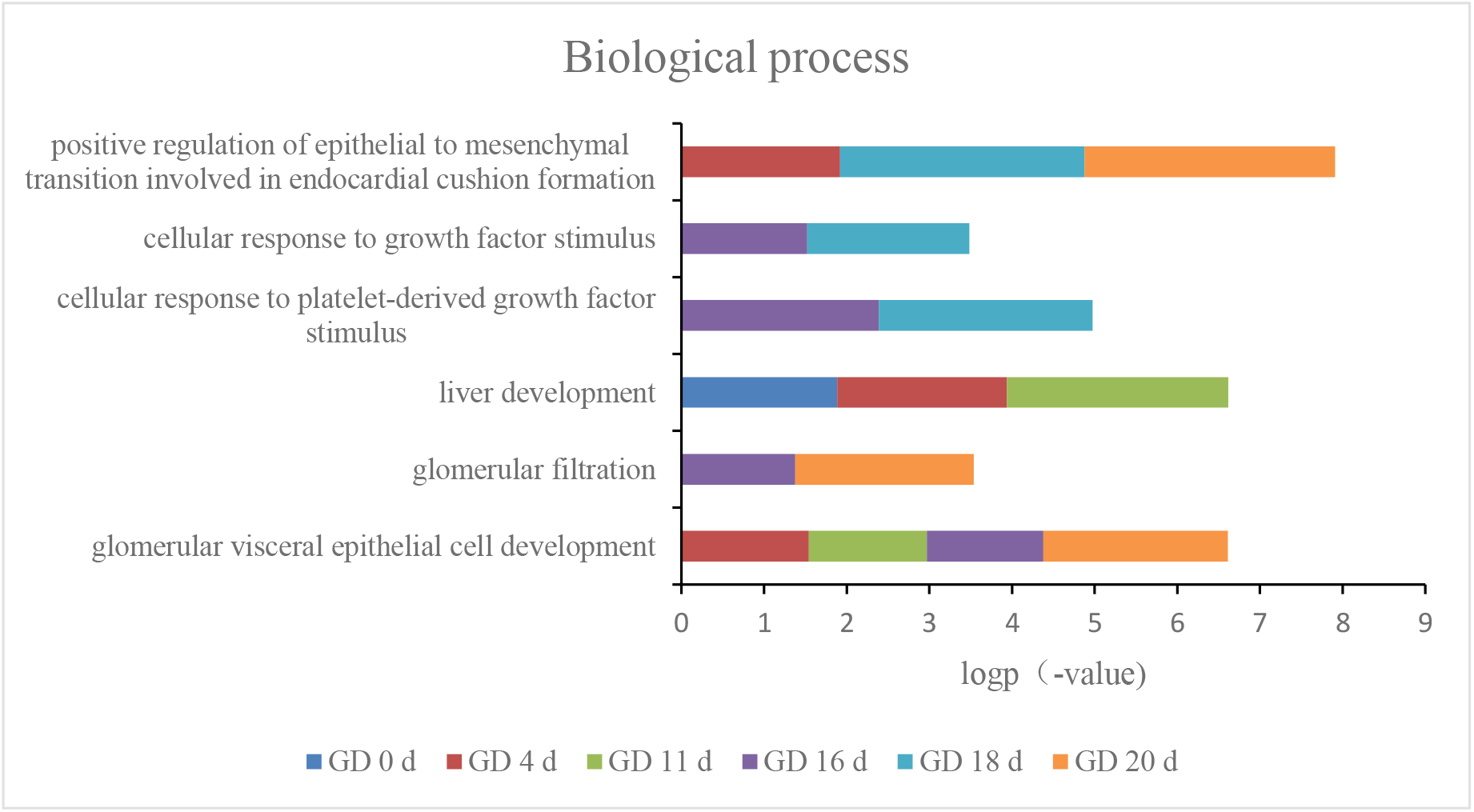
Changes in pathways related to organ development over time

### 3.5 Functional analysis of the differentially expressed proteins in the pregnant group related to the maternal coagulation system

From the first to the third trimester, with increasing gestational age, coagulation function is gradually enhanced, including increased thrombin generation and fibrinolysis. These two factors co-occur and, in most cases, can lead to a hypercoagulable state. These changes are thought to maintain placental function during pregnancy and prevent massive blood loss during delivery (34).

Figure 8A shows the biological pathways on GD 20 d, with the extrinsic prothrombin activation pathway, GP6 signaling pathway, coagulation system, intrinsic prothrombin activation pathway, and prostanoid biosynthesis being related to coagulation function (Fig. 8A). The interaction of platelets with collagen via the GP6 receptor leads to platelet activation and adhesion, a process required for thrombosis. GP6 is a platelet transmembrane glycoprotein that plays an important role in collagen-initiated signal transduction and platelet procoagulant activity (35). Prostanoid biosynthesis and endogenous prostaglandin production are associated with labor in humans, prostaglandin levels continue to rise during labor, and prostaglandins appear to be involved in the increased contractile activity observed during labor (36). The changes in these biological processes related to procoagulation are shown in Fig. 8B. The trend of coagulation biological process between GD 14 d and GD 20 d showed that the pathway was significantly upregulated on GD 20 d. In addition, compared with GD 14 d, the number of coagulation pathways also increased significantly on GD 20 d, suggesting that the coagulation function of pregnant mice continued to increase before delivery.

**Fig. 8A.**
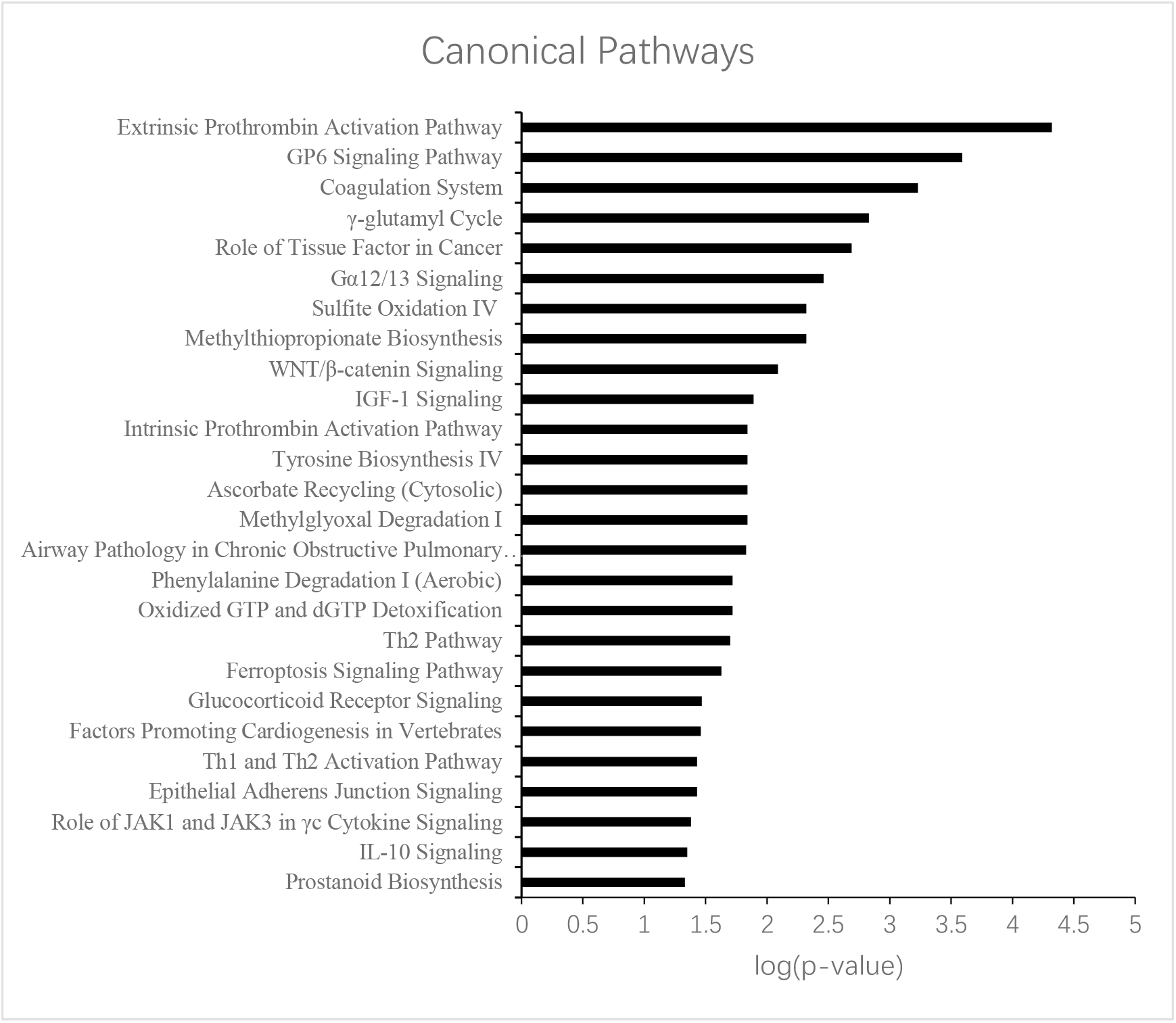
Analysis of biological processes by IPA (P*<0.5) at GD 20 d

**Fig. 8B.**
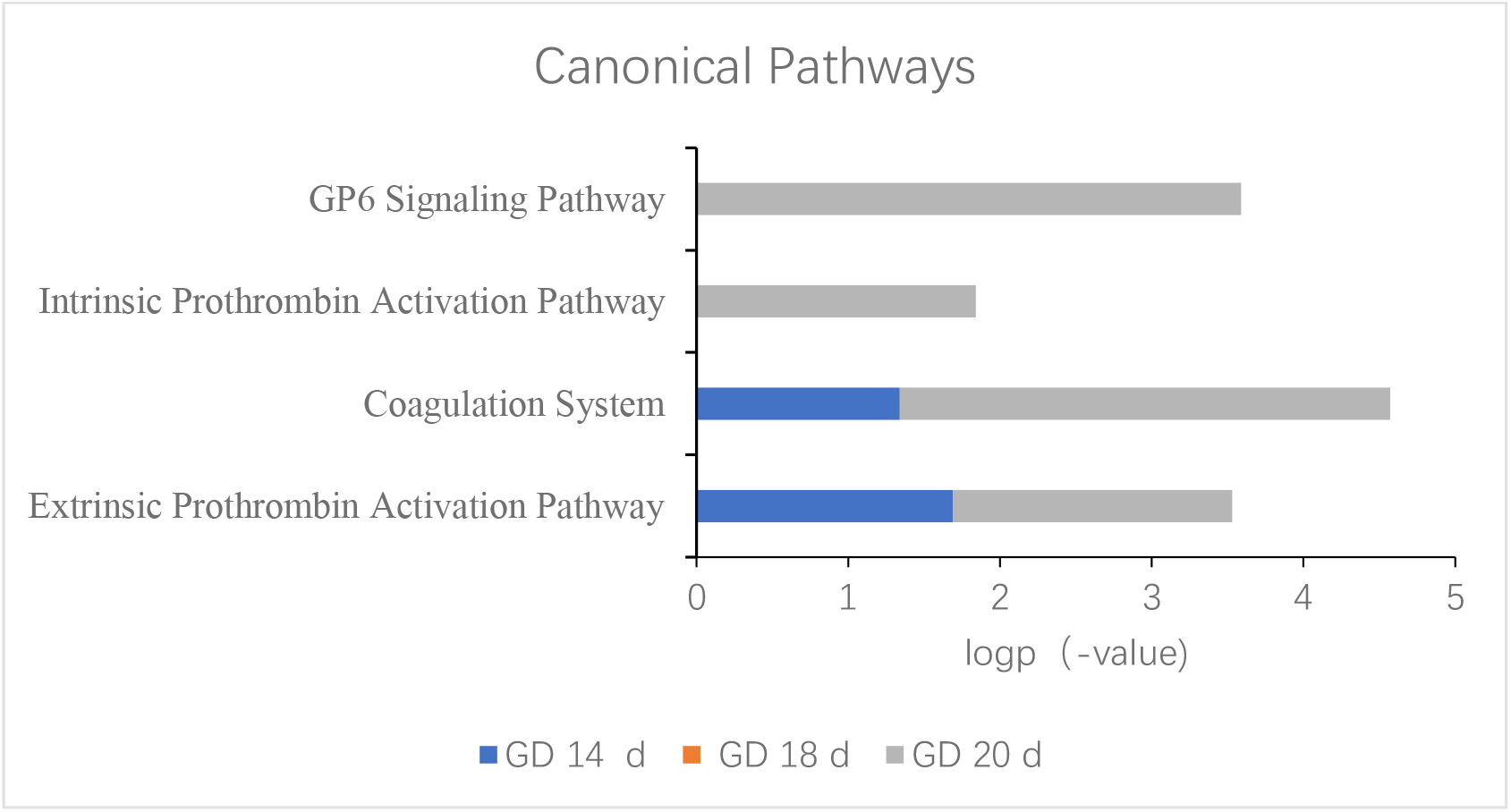
Changes in coagulation-related pathways on GD 20 d Fig. 9A The line plot for the relative abundance of coagulation-related protein MOR8A3 during pregnancy

**Fig. 8C.**
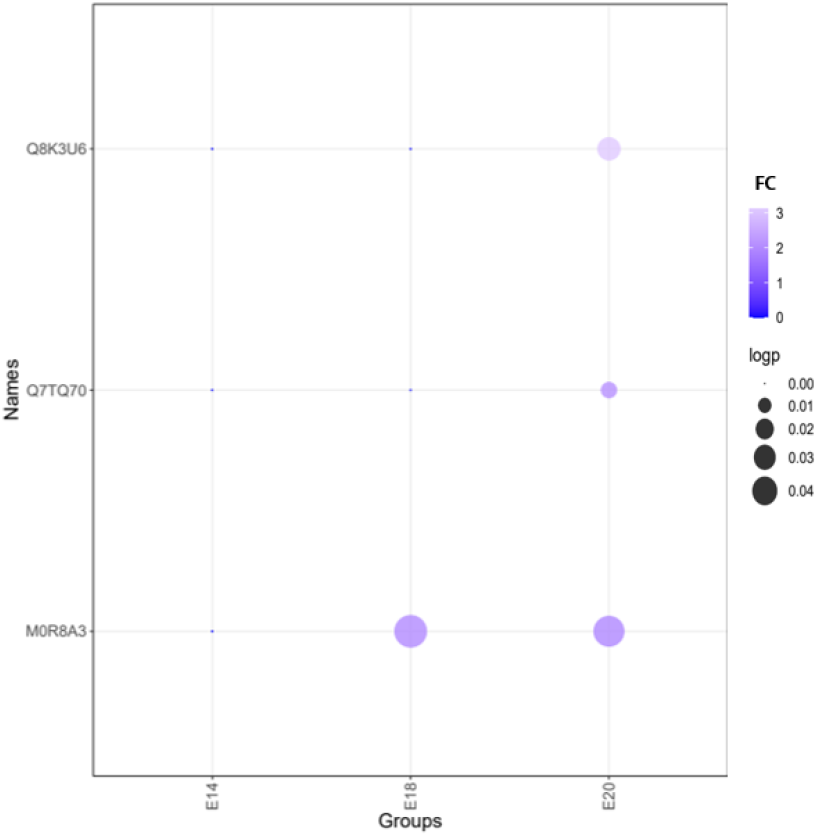
Change trends of coagulation-related proteins on GD 20 d.

To assess changes in coagulation-related proteins, their relative abundances were analyzed during pregnancy (Fig. 9A). MOR8A3, Glycoprotein 6 (platelet), was expressed at low levels in early pregnancy, but up-regulated at GD 14 d and maintained at high levels before delivery (GD 20 d). Q7TQ70, Fibrinogen alpha chain, this protein is an important component in the formation of insoluble fibrin matrix, fibrin is one of the main components of blood clots and plays an important role in hemostasis. This protein was also up-regulated at GD 14 d and remained higher than the first trimester until delivery. Similarly, the procoagulant-related proteins (Fig. 9B) fold change became larger, and P value was more significant over time. Changes in the coagulation system revealed by the urinary proteome were consistent with reported changes before labor.

**Fig. 9A.**
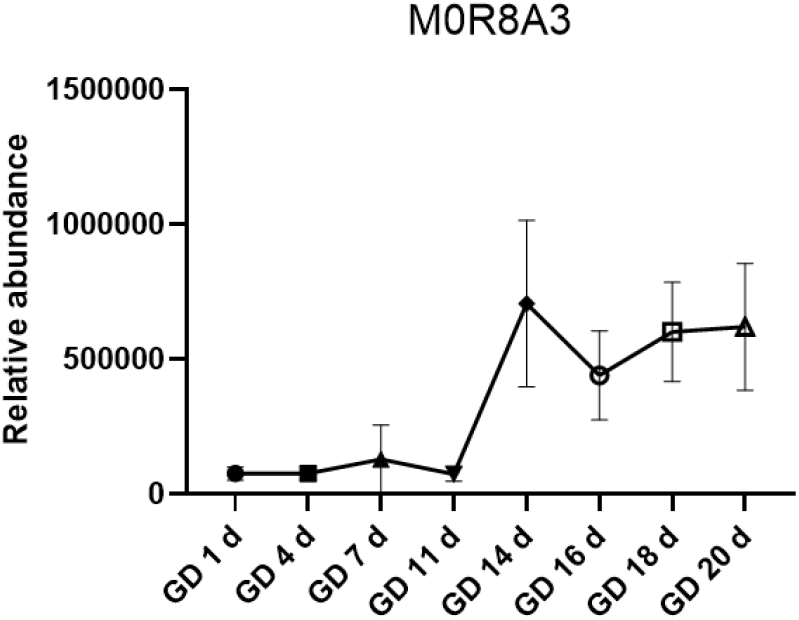
The line plot for the relative abundance of coagulation-related protein MOR8A3 during preganancy

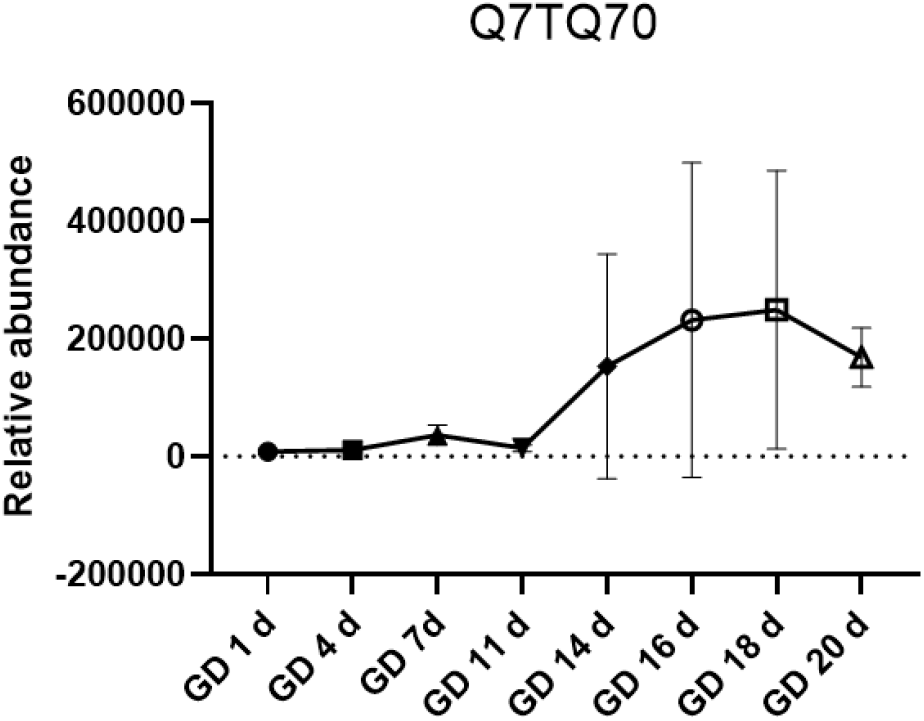
The line plot for the relative abundance of coagulation-related protein Q7TQ70 during pregnancy

Our results show that the urinary proteome can reflect changes in different gestational stages. During the period from fertilization to implantation in rats, GD 1 d, GD 4 d and GD 7 d were enriched in many pathways related to embryo implantation and trophoblast differentiation. During the second and third trimesters, pathways related to the development of embryonic organs, such as the development of the heart, heart looping and endocardial cushion formation, were consistent with the developmental times reported in previous studies. The timing of the emergence of alveolar development-related pathways is in good agreement with previous studies. The development of pancreatic islet B cells also occurs during the main proliferation and differentiation periods of pancreatic cells. In addition, some other tissues and organs developed during this period, such as the liver and glomerulus. All of these pathways were manifested only in the pregnancy group and not in the control group. On the 20 day of gestation, many coagulation-related pathways were enriched, and the significance of coagulation-related proteins and pathways continued to increase, which was consistent with the change trend of prenatal coagulation function reported in the literature. In addition, the related proteins of embryonic organ development began to work at a very early stage (GD 0 d), indicating that urine may reflect the embryonic development status at an early stage. Based on these results, the urinary proteome is very sensitive in reflecting the process of different stages of pregnancy, providing information on normal pregnancy and embryonic growth and development.

Although this study is based on animal experiments, the results suggest that more clinical samples need to be included to explore the pregnancy process reflected by the human urine proteome in the future. The normal urine proteome of pregnant women may serve as a clinical database and be used as an effective method for the obstetric detection of abnormal pregnancy.

## 4 Conclusion

In summary, the urine proteome can reflect the process of normal rat pregnancy, including maternal changes and embryonic development, laying a foundation for further clinical research. The urine proteome has the potential to be a new and effective method for embryonic status determination

## Acknowledge

The datasets presented in this study can be found in online repositories: http://www.proteomexchange.org/. The database can be found since publication.

